# Intratumoral plasma cells mediate CD8+ T cell infiltration and successful immune checkpoint blockade therapy in *de novo* MPNSTs

**DOI:** 10.64898/2026.02.18.706680

**Authors:** Joshua J. Lingo, Ryan Reis, Chantal Allamargot, Juan A. Raygoza Garay, Courtney A. Kaemmer, Elizabeth C. Elias, Ellen Voigt, Ali Jabbari, Connor R. Wilhelm, Alexander W. Boyden, Nitin J. Karandikar, Patrick Breheny, David K. Meyerholz, Rebecca D. Dodd, Jon C.D. Houtman, Benjamin W. Darbro, Dawn E. Quelle

**Affiliations:** Cancer Biology Graduate Program, University of Iowa, Iowa City, Iowa; The Department of Neuroscience and Pharmacology, University of Iowa, Iowa City, Iowa; Medical Scientist Training Program, University of Iowa, Iowa City, Iowa; Central Microscopy Core Research Facility, University of Iowa, Iowa City, Iowa; Holden Comprehensive Cancer Center, University of Iowa, Iowa City, Iowa; Maximizing Access to Research Careers (MARC) Program, University of Iowa, Iowa City, Iowa; The Department of Dermatology, University of Iowa, Iowa City, Iowa; The Department of Pathology, University of Iowa, Iowa City, Iowa; The Department of Biostatistics, University of Iowa, Iowa City, Iowa; The Department of Internal Medicine, University of Iowa, Iowa City, Iowa; The Department of Microbiology and Immunology, University of Iowa, Iowa City, Iowa; The Department of Pediatrics, University of Iowa, Iowa City, Iowa

**Keywords:** MPNST, plasma cells, PD-L1, CD8+ T cells, immune checkpoint blockade (ICB)

## Abstract

**Background:** The role of intratumoral plasma cells in immune checkpoint blockade (ICB) therapy has never been tested although their presence is linked with improved patient response and survival. Malignant peripheral nerve sheath tumors (MPNSTs) are deadly sarcomas with minimal responsiveness to ICB therapies. Strikingly, drugs inhibiting cyclin-dependent kinases 4/6 (CDK4/6) and MEK sensitize *de novo* MPNSTs to immunotherapy targeting programmed death-ligand 1 (PD-L1), which correlates with increased intratumoral plasma cells. Here, we tested if plasma cells mediate the MPNST response to anti-PD-L1 therapy.

**Methods:** Anti-tumor activity of PD-L1 inhibition, with or without CDK4/6-MEK inhibition, was measured in *de novo* MPNSTs within wild-type versus plasma cell-deficient mice. Plasma cell-dependent effects of CDK4/6-MEK inhibition on priming the MPNST immune environment were determined by single cell transcriptomics and immunostaining.

**Findings:** MPNSTs lacking plasma cells failed to respond to anti-PD-L1 monotherapy and were no longer sensitized to immunotherapy by CDK4/6-MEK inhibition. Plasma cell-deficient MPNSTs exposed to CDK4/6-MEK inhibitors had impaired antigen presentation on major histocompatibility class I (MHC-I) and decreased CD8+ T cell infiltration and activation. Complementary analyses of human sarcomas showed increased intratumoral plasma cell signatures prognose better patient survival.

**Interpretation:** Plasma cells favorably remodel the tumor immune environment by increasing CD8+ T cell infiltration and are critical for successful ICB therapy in MPNSTs. This work may help inform ICB treatment strategies and cancer patient stratification for many different tumor types.

**Funding:** This research was supported by University of Iowa Sarcoma Research Program awards and NIH grants T34-GM141143, T32-GM067795, F31-CA281312, P30-CA086862, and R01-NS119322.

**Research in Context:** *Evidence before this study:* For many types of cancer, intratumoral plasma cells have been *correlated* with better patient survival and improved response to immune checkpoint blockade (ICB) therapies. However, the biology underlying those associations *is not understood* and no study has examined the requirement of plasma cells in immunotherapy response. Compelling data in malignant peripheral nerve sheath tumors (MPNSTs) showed that dual kinase inhibition of oncogenic CDK4/6 and MEK induced intratumoral plasma cell accumulation and sensitized tumors to ICB therapy. While CDK4/6-MEK inhibition is known to enhance antitumor immunity in other tumor types by CD8+ T cells or natural killer (NK) cells, a role for plasma cells has never been explored.

*Added value of this study:* Studies were performed in MPNSTs, an under-researched cancer that normally responds poorly to ICB monotherapies. This is the first investigation to show that intratumoral plasma cells are essential for successful ICB therapy and they support anti-tumor immunity by promoting a pro-inflammatory, CD8+ T cell state involving MHC-I antigen presentation. Findings provide new insight into immunomodulatory effects of CDK4/6-MEK inhibitor therapies, revealing plasma cells are needed for those drugs to activate CD8+ T cell mediated antitumor immunity.

*Implications of all the available evidence:* The fundamental advance in understanding how plasma cells promote successful ICB immunotherapy is likely applicable to other solid tumors and may guide novel therapeutic strategies in which plasma cell-inducing agents are combined with ICB antibodies. Moreover, an increased presence of intratumoral plasma cells in tumor specimens may streamline clinical decisions regarding which patients are most likely to benefit from ICB therapy.

## Introduction

Malignant Peripheral Nerve Sheath Tumors (MPNSTs) are an aggressive subtype of sarcoma that originates from the myelinating nerve sheath. MPNSTs can arise sporadically or in patients with Neurofibromatosis Type 1 (NF1), the most common, heritable tumor predisposition syndrome worldwide ^1,2^. Unfortunately, MPNSTs are notoriously unresponsive to chemotherapy, lack approved targeted therapies, and generally are insensitive to immune checkpoint blockade (ICB) therapy. The latter is thought to reflect an immunologically “cold” nature of MPNSTs although it is an understudied area in MPNST research ^3^.

MPNSTs are driven by hyperactivated Ras/MEK/ERK signaling initiated by loss of *NF1*, a tumor suppressor gene that encodes neurofibromin, a Ras GTPase activating protein (GAP) that converts Ras into its inactive, GDP-bound form ^4^. In NF1 individuals, the mutation of *NF1* promotes formation of benign plexiform neurofibromas (PNFs), precursor lesions that can transform into MPNSTs in up to one-third of patients ^2^. Additional alterations cooperate with *NF1* loss to support PNF transition into MPNSTs ^2,5-7^. Two of the most common changes driving that process are loss of *INK4a* and *ARF* (collectively called *CDKN2A*) ^7^ and overexpression of RABL6A ^8^, events that converge functionally to activate oncogenic cyclin-dependent kinases 4 and 6 (CDK4/6) and turn off the Retinoblastoma 1 (RB1) tumor suppressor ^9,10^. Elevated Ras/MEK activity also upregulates CDK4/6 through MYC-mediated transcription of *CDK4, CDK6*, and *CCND1* (which encodes the cyclin D1 regulatory subunit required for CDK4/6 activity) ^11,12^. Efforts targeting MEK or CDK4/6 individually in preclinical MPNST models have yielded unsatisfactory results due to rapid resistance ^13-19^, prompting exploration of combination therapies targeting those factors in MPNSTs.

Preclinical studies pairing CDK4/6 and MEK inhibitors have proven effective at suppressing Ras pathway-driven cancers through activation of anti-tumor immunity. This includes MPNSTs ^20^, ovarian cancer ^21^, colorectal cancer ^22^, melanoma ^23,24^, as well as adenocarcinomas of the lung ^25^ and pancreas ^26^. In *de novo* MPNSTs induced by *Nf1/Ink4a/Arf* inactivation in the mouse sciatic nerve, dual CDK4/6-MEK inhibition provokes an anti-tumor immune response featuring activated CD8+ T cells and intratumoral plasma cell accumulation ^20^. The plasma cell phenotype, the first to be reported for this drug combination in any tumor, is compelling because intratumoral plasma cells predict prolonged patient survival ^27-30^ and improved response to ICBs ^28,31-34^ for most human cancers. Indeed, combining CDK4/6 and MEK inhibition with ICB therapy targeting programmed death ligand 1 (PD-L1) causes sustained MPNST regression and extended mouse survival with some mice exhibiting cure ^20^. Synergistic tumor killing by combining CDK4/6 and MEK inhibitors with another ICB agent, anti-PD-1 antibodies, has been achieved in other tumor types ^21,25,26^. However, no study has investigated or determined the requirement of plasma cells for mediating the anti-tumor immune response to ICB therapy.

Here, we directly test the role of plasma cells in the response to anti-PD-L1 therapy in *de novo* MPNSTs in mice that either contain or lack plasma cells. Our findings demonstrate that plasma cells mediate a CD8+ T cell, pro-inflammatory state following CDK4/6-MEK inhibition that is critical for successful ICB therapy. Understanding the unique role of plasma cells in controlling therapy-induced, anti-tumor immunity may have broad relevance to the effective treatment of solid cancers with immune-based therapies.

## Methods

### Sex as a biological variable

Our study examined male and female animals, and similar findings are reported for both sexes.

### Primary MPNST generation

Mice were housed in barrier rooms with a 12-hour light-dark cycle and free access to food and water. All mouse handling was conducted in compliance with the Institutional Animal Care and Use Committee (IACUC) policies at the University of Iowa. These requirements adhere to the NIH Guide for the Care and Use of Laboratory Animals and the Public Health Service Policy on the Humane Care and Use of Laboratory Animals. All efforts were made to minimize animal suffering. Primary MPNSTs were generated via adenoviral delivery of Cas9 and sgRNAs targeting *Nf1* and *Cdkn2a* to the left sciatic nerve as previously reported ^35^. Tumors were initiated in wild-type C57BL/6N and plasma cell-knockout (PC-ko, *AID*−/−; *µS*−/−, Activation-induced cystine deaminase (AID)/secretory mu-chain (µS) double knockout) mice ^36^. Male and female mice were used equally.

### Flow cytometric validation of *AID*−/−; *µS*−/− mice

Fresh splenocytes were harvested from wild-type and PC-ko mice via physical disruption of the spleen and then passed through a 70µM strainer ^37^. Single cell suspensions of splenocytes were stained for viability with Zombie NIR (Biolegend, cat. 423105) and Fc receptors were blocked with anti-CD16/32 (Biolegend, clone 93) for 10 minutes on ice in the dark. Samples were washed in Cell Staining Buffer (Biolegend, 420201) and incubated with the following antibodies for 30 minutes on ice: CD45 (clone 30-F11, BV510, Biolegend #103137), B220 (clone RA3-RB2, APC/Fire 810, Biolegend #103277), CD3 (clone 17A2, Spark NIR 685, Biolegend #100261), and CD138 (clone 281-2, APC, Biolegend 142505). Splenocytes were washed via centrifugation, fixed in FluoroFix Buffer (Biolegend, cat. 422101), washed again, and stored overnight at 4C. Samples were analyzed on the Cytek Aurora. Flow cytometric analyses were performed using FlowJo 10.9.0 (Becton, Dickson and Company). Plasma cells were identified as live, CD45+CD3-B220-CD138+ singlets. Fluorescence minus one (FMO) controls were used to identify positive and negative populations.

### *In vivo* tumor growth analyses and drug treatment conditions

Tumor size was measured with calipers and tumor volume was calculated using the formula (length x width x thickness x π) / 6. After tumor detection (∼250 mm3), mice were randomly enrolled into treatment groups. Daily, mice received, by oral gavage, either a vehicle control (50 mM sodium-L-lactate, pH 4.0 plus 1% DMSO) or kinase inhibitors (100 mg/kg palbociclib [CDK4/6 inhibitor] plus 1 mg/kg mirdametinib [MEK inhibitor]). For the first three weeks of therapy, starting upon tumor detection, mice received IgG control (10 mg/kg anti-keyhole limpet hemocyanin [BioXCell BE009]) diluted in InVivoPure pH 7 [BioXCell IP0070]) or anti-PD-L1 (10 mg/kg anti-PD-L1 [BioXCell BE0101] diluted in InVivo Pure pH 6.5 [BioXCell IP0065]) intraperitoneally, twice per week for a total of 6 doses. Each mouse was euthanized after the tumor reached ∼1250 mm^3^. Tumors were immediately harvested and sectioned for RNA/protein analysis (flash-frozen) and histopathological analysis (formalin-fixed paraffin-embedded, FFPE).

### Preparation of *de novo* MPNSTs for single cell RNA sequencing

After tumor detection (∼450 mm^3^), wild-type C57BL/6N and *AID*−/−; *µS*−/− mice received 100 mg/kg palbociclib plus 1 mg/kg mirdametinib daily by oral gavage until they regressed to ∼250 mm^3^, which occurred within 4-6 days. Upon reaching the desired endpoint, mice were euthanized and tumors were immediately excised. Each tumor was divided into sections for flash freezing, immunohistochemistry (FFPE), or digested into single cell suspensions for single cell RNA sequencing (scRNA seq). To develop single cell suspension, small portions of each tumor were cut into small cubes (∼8 mm^3^). Minced tumors were then dissociated using the Mouse Tumor Dissociation Kit (Miltenyi Biotec 130-096-730) with all dissociation steps requiring media performed in DMEM. Tumor dissociation was completed using the gentleMACS Dissociator (Miltenyi Biotec cat. #130-093-235). After mechanical dissociation, samples were enzymatically digested according to manufacturer protocols on an orbital shaker at 250 rpm at 37C for 45 minutes. Suspensions were filtered using 70 µM mesh strainer and percent viability determined via trypan blue exclusion. Viable suspensions were cryopreserved in 90% fetal bovine serum (FBS) and 10% DMSO at a concentration of 1 ×10^6^ cells/mL. Cryopreserved single cell suspensions of *de novo* MPNSTs were thawed at 37× C on the day of sequencing and viability reconfirmed via trypan blue exclusion. Only single cell suspensions with viability greater than 80% were included in GEM preparation using the 10X Genomics Chromium controller at the Iowa Institute of Human Genetics Genomic Division. cDNA libraries were generated using the Chromium Single Cell 3’ v3.1 kit. Sequencing was conducted on the Illumina NovaSeq 6000 using an S1 flow cell.

### scRNA sequencing analyses

FASTQ files were concatenated across sequencing lanes and aligned to the GRCm39-2024-A mouse reference genome using cellranger (v7.0.1) with default settings. Ambient RNA was removed from each raw_feature_bc_matrix.h5 file via cellbender (v0.3.0), using default parameters except for the following: epochs = 50; this choice was made following inspecting of the ELBO convergence plots for each sample. The resulting filtered.h5 outputs were loaded into Rstudio (R v4.4.1) and converted into Seurat objects using the Seurat (v5.1.0) package with min.cells = 5 and min.features = 300. Percent mitochondrial and ribosomal reads were calculated for each cell and doublets called using scDblFinder (v1.18.0). One sample (B_009, WT) was excluded due to what appeared to be persistent RNA contamination (ambient RNA), and the remaining samples were merged into a single Seurat object. Additional filtering was then applied via the following: db.class == “singlet” & nFeature_RNA > 250 & nCount_RNA > 400 & nCount_RNA < 150000 & percent.mt < 25 & percent.rb < 40. The filtered dataset underwent normalization with SCTransform, regressing out percent.mt and nCount_RNA. Principal component analysis was performed on the SCT assay, followed by harmony (v1.2.0) integration to correct for batch effects. Uniform Manifold Approximation and Projection (UMAP) embeddings were computed on the harmony-corrected dimensions, and graph□based clustering was completed using the first 20 dimensions. Marker genes were identified with the Seurat FindAllMarkers function, and major cell□type identities were assigned based on these results.

CellChat (v2.1.2) was used to identify differences in cell-cell communication between the genotypes of interest. CellChat objects were generated separately for each genotype from their SCT assays, merged into a combined CellChat object, and a comparative analysis was performed, comparing the PC_KO to the WT genotype results. Metaprograms (MPs) were identified using GeneNMF (v0.6.0). MPNST cells were subset and split by sample. The SCT assay was used as input to multiNMF, with parameters k = 6:8, L1 = c(0,0), scale = TRUE, center = TRUE, and nfeatures = 1,000. Eight consensus MPs were identified via getMetaPrograms (max.genes = 50) and visualized in a heatmap of overlapping genes. Individual cells were scored for expression of each MP using AddModuleScore_UCell. Gene ontology was performed for each MP using clusterProfiler. For major cell types of interest, these cells were subset from the original object and re-processed individually to create celltype-specific pca and umap reduced dimensions and clusters. Marker genes were again identified with FindAllMarkers, and the clusters were labeled according to these differentially expressed genes (DEGs). DEGs were also identified as a function of genotype for genotype-specific gene onotology. Heatmaps were generated via the R2easyviz (v0.1.0) package. Cells were down sampled and grouped according to their subcluster and the expression of the top DEGs, as a function of percent difference in expression (pct.1 - pct.2), for each was scaled and centered for visualization. Stacked bar charts were generated for each subcluster of interest and colored by genotype. The proportion of cells from each genotype was calculated as a function of total cells within each cluster and presented as percent of total. Select feature plots and dot plots were generated using the Seurat package.

### Mouse tumor RNA isolation and quantitative RT-PCR analyses

RNA was isolated from flash frozen mouse tumors using Qiagen RNAeasy Mini Plus Kit. RNA yield and purity were obtained using the Trinean DropSense 16. cDNA was synthesized from 200 ng total RNA using SuperScriptTM III First-Strand Synthesis System (Invitrogen, cat. 18080051). Diluted cDNA was used for real-time PCR (cycle conditions: denaturation at 95C for 10 minutes, followed by 40 cycles of 95C for 15 seconds and 45-50C for 1 minute, using SYBR Green for detection (Bio Rad, cat. 1708880). Gene-specific primers were used to determine the fold change in mRNA expression using the 2-ΔΔCt method, calibrated to *Gapdh* mRNA expression. Primer sequences are in (**Supplementary Table 1**).

### Histopathological analysis of mouse MPNSTs

Upon euthanasia, tumors were harvested and stored in 10% neutral-buffered formalin. Tumors were embedded in paraffin, and sectioned (5µm) onto glass slides, and hydrated. Heat-induced epitope retrieval with Citrate-based Antigen Unmasking Solution pH 6.0 (Vector Laboratories H-3300-250) at greater than 95C for 5 minutes followed by 5 minutes of rest. This was repeated once. After blocking, tissue sections were immunostained with the antibodies in **Supplementary Table 2** in blocking buffer (5% Normal Donkey (Sigma, D9663)/Goat Serum (Sigma, G-9023) in PBS or Vector M.O.M. Immunodetection kit (Vector Laboratories, BMK-2202) overnight at 4C. Sections were then washed in PBS and incubated with goat anti-rabbit IgG F(ab’)_2_ conjugated to Alexa-Fluor 488, 568, or 647 for one hour at room temperature. All secondary antibodies were used at a 1:500 dilution in blocking buffer. Antibodies that required signal amplification (SA) for visualization were incubated with biotinylated goat antibodies raised against the host species of the primary antibody for 1 hour followed by a PBS wash and incubation with streptavidin-Alexa Fluor 647 conjugate for one hour. Before any work with biotinylated antibodies, endogenous biotin was blocked with egg substitute solution and fat-free milk for 15 minutes each. After all antibody incubations, slides were washed with PBS and glass coverslips were mounted using Prolong Glass Antifade Mountant with NucBlue (ThermoFisher P36981). Sections of tissue were examined by at least two observers (including a pathologist) and morphometric analysis was performed using the post-examination method of masking to group assignment ^38^. At least 4 high-power field images (HPF, 40X) were obtained for each tumor section and stain using a Zeiss LSM 980. Positive cells per HPF were counted manually and fluorescent intensity (relative fluorescent units, RFU) was quantified using Fiji/ImageJ. Lymphoid infiltration was quantified from hematoxylin and eosin-stained tissue sections. Percent lymphoid area was defined as the total area of lymphoid infiltration divided by the total area of the tumor section.

### Analysis of TCGA bulk RNA sequencing

Immune cell prediction was performed on publicly available RNA sequencing data from TCGA portal (SARC cohort). CIBERSORT was run using the LM22 signature matrix file, in relative (not absolute) mode, with quantile normalization disabled, and 1000 permutations. Intercellular correlations were determined the Pearson correlation method. High and low cell fractions were determined by the upper and lower 45^th^ percentile, respectively, after the exclusion of any samples with null values. Kaplan-Meier analysis was performed to determine statistical differences in cohort survival. Detailed information for number at risk for each time point are available in **Supplementary Table 3**.

### Statistics

Quantified data are presented as the mean and all error bars are SEM. P values, unless otherwise specified, were obtained by One-way ANOVA and adjusted for multiple comparisons using the indicated method. P values less than 0.05 were considered statistically significant. Statistical analysis of median survival was performed using Kaplan-Meier Analysis.

### Study approval

All mouse studies adhered to protocols approved by the Institutional Animal Care and Use Committees at the University of Iowa (Protocol 3092074: Targeting RB1 Pathway in Sarcomas). All efforts were made to minimize animal suffering.

## Results

### Plasma cells are essential for MPNST response to anti-PD-L1 therapy

It is increasingly appreciated that the presence of B cells and plasma cells in human tumors correlates with improved response to ICB therapy ^28,31-34^, although their exact role in anti-tumor immunity is poorly understood. Preclinical studies in melanoma and ovarian cancer showed that B cells are required for response to ICB therapy alone or combined with other therapies, including CDK4/6 inhibitors ^21,39,40^. No study, however, has investigated the role of plasma cells in ICB therapy response.

Given our recent findings that dual CDK4/6-MEK inhibition in de novo MPNSTs synergistically increases intratumoral plasma cells (not B cells) and sensitizes tumors to anti-PD-L1 therapy ^20^, we sought to test the requirement of plasma cells for ICB response in MPNSTs. To do so, *de novo* MPNSTs were generated in wild-type (WT) mice as well as *AID−/−; µS−/−* knockout mice devoid of plasma cells. The loss of activation-induced cystine deaminase (*AID*) and the secretory mu-chain (*µS*) genes (hereafter termed PC-ko for plasma cell knockout) yields agammaglobulinemic mice that retain B cells and plasmablasts but selectively lack plasma cells ^36^. The T cell compartment in PC-ko mice is intact ^36^. Using established methods ^8,20,35,41^, *de novo* MPNSTs were induced by CRISPR-Cas9 editing of endogenous *Nf1* and *Cdkn2a* (encodes both *Ink4a* and *Arf*) in the sciatic nerve of wild-type and PC-ko mice (Figure 1a). MPNSTs in PC-ko mice initiated similarly to those in wild-type mice with no difference in tumor onset (Figure 1b). The loss of plasma cells in PC-ko mice was validated by flow cytometric identification of live splenic CD45+B220-CD138+ cells ^36^ (Figure 1c). These data show that plasma cells do not influence MPNST initiation.

**Figure 1.**
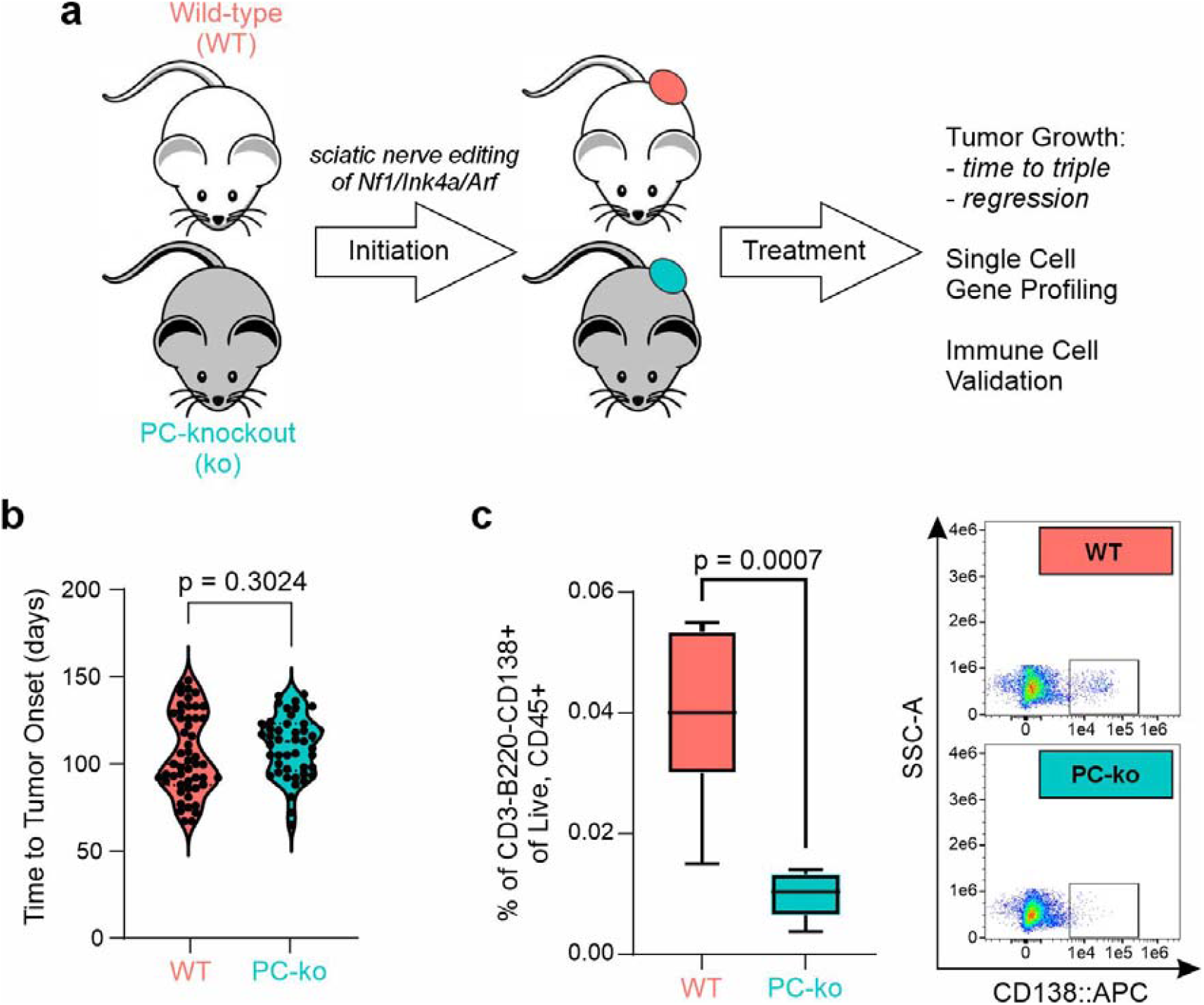
Loss of plasma cells does not alter the initiation of *de novo* MPNSTs generated by inactivation of *Nf1* and *Cdkn2a*. a) Schematic depicting generation of *de novo* MPNSTs via CRISPR-mediated inactivation of *Nf1* and *Cdkn2a* (encoding *Ink4a* and *Arf*) in the sciatic nerve of wild-type (WT) and AID/µs−/− mice (PC-knockout, PC-ko). b) Quantification and comparison of time to tumor onset of all MPNSTs in this study. c) Population frequencies of plasma cells relative to all Live, CD45+ cells, with representative pseudocolor plots on the right. Plasma cells were defined as CD3-B220-CD138+ cells. P values were obtained using a Student’s t test.

To directly determine the necessity of plasma cells for tumor response to ICB therapy, *de novo* MPNSTs in wild-type and PC-ko mice were treated with anti-PD-L1 antibodies alone, inhibitors to CDK4/6 (palbociclib) plus MEK (mirdametinib), or the triple combination versus vehicle with IgG control. Therapy began once tumors reached approximately 250-350 mm^3^ in size. Tumors in wild-type mice responded to the treatments in an identical manner to our previous study ^20^. Control vehicle/IgG-treated tumors tripled in volume at similar rates in wild-type and PC-ko mice (Figure 2a). Likewise, in both genotypes, treatment with CDK4/6-MEK inhibitors similarly delayed MPNST growth relative to controls. Differences in tumor progression between wild-type and PC-ko mice were only seen following treatment with anti-PD-L1 therapy, either alone (vehicle + anti-PDL1) or when combined with CDK4/6-MEK inhibition (CDK4/6i-MEKi + anti-PDL1). Under those conditions, the slower tumor growth caused by anti-PD-L1 therapy in wild-type animals was completely lost in PC-ko mice (Figure 2a). Thus, plasma cells are required for MPNST response to ICB therapy.

**Figure 2.**
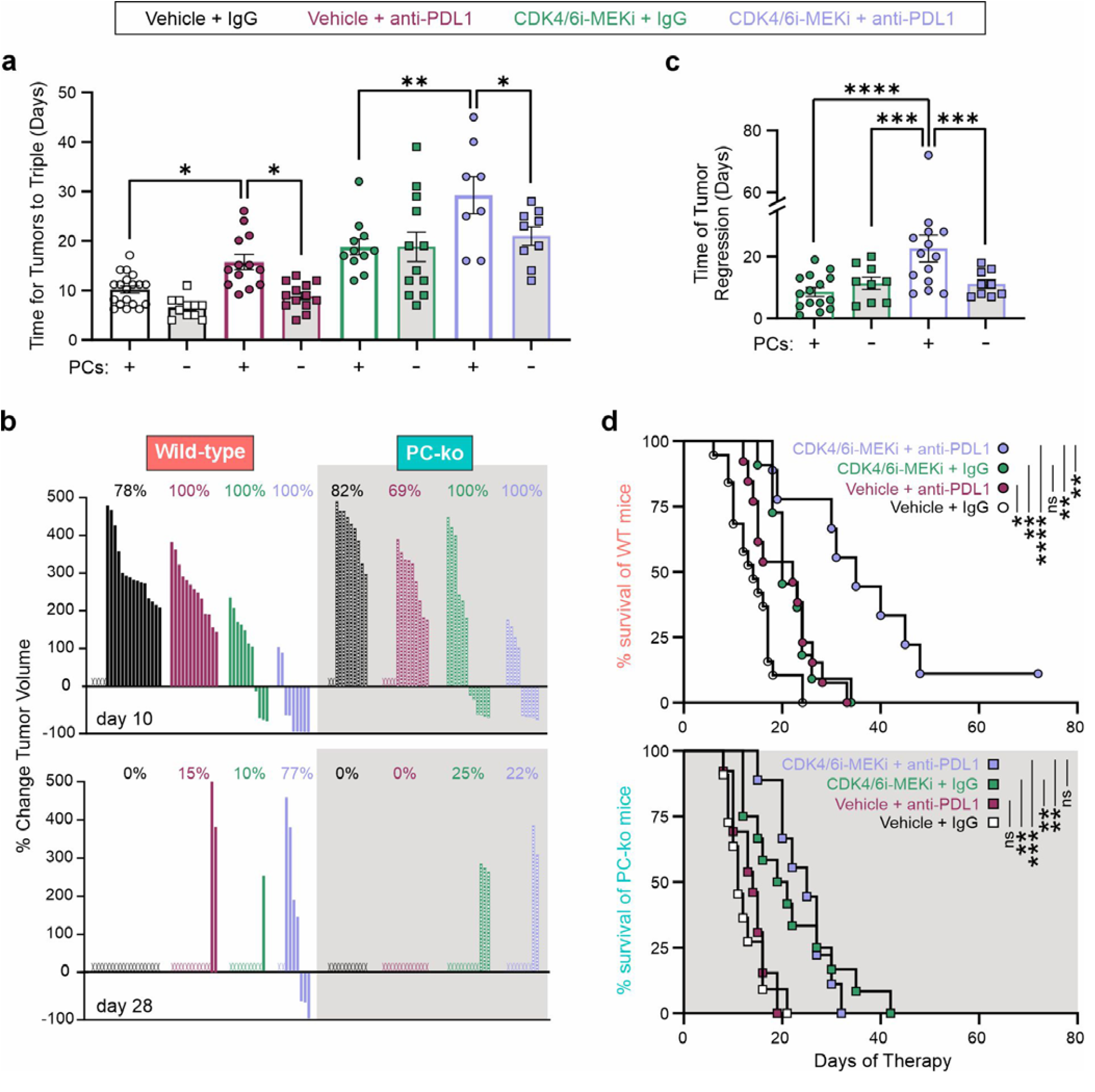
Plasma cells are required for successful response to anti-PD-L1 therapy in de novo MPSNTs. Wild-type and PC-ko mice bearing *de novo* MPNSTs, initiated by *Nf1/Ink4a/Arf* editing in the sciatic nerve, were treated daily with vehicle or CDK4/6-MEK inhibitors (palbociclib at 100 mg/kg, mirdametinib at 1 mg/kg). Mice also received 2 weekly i.p. injections of IgG control or anti-PD-L1 antibodies for the first 3 weeks of therapy. a) Time (in days) it took for MPNSTs to triple in volume. b) Waterfall plots showing percent change in tumor volume at days 10 (top) and 28 (bottom) of treatment. c) Time (in days) that tumors treated with CDK4/6 and MEK inhibitors +/- anti-PD-L1 therapy regressed, as defined by the measured tumor volume being less than volume at detection. d) Survival of wild-type (WT, top) and plasma cell-deficient (PC-ko, bottom) mice bearing MPNSTs treated with the indicated therapies. Error bars are SEM. P value was obtained via two-way ANOVA (panels a and c) or through Kaplan-Meier analysis (panel d). ns, not significant. In panels a and c, only statistically significant comparisons are shown (*, P < 0.05, **, P, < 0.01, ***, P < 0.001, ****, P < 0.0001).

The above results corresponded with more pronounced tumor regression in plasma cell-containing MPNSTs treated with CDK4/6-MEK-PDL1 targeted therapy. A greater magnitude of tumor shrinkage (seen by the negative fold change in tumor volume) was sustained over time for CDK4/6-MEK-PDL1 targeted MPNSTs in wild-type mice, but this was lost in PC-ko mice (Figure 2b). Indeed, wild-type tumors treated with CDK4/6-MEK-PDL1 inhibitor therapy had a >2-fold longer average time of tumor regression relative to identically treated PC-ko tumors, which instead matched the shorter time of tumor regression caused by CDK4/6-MEK inhibitors alone in wild-type and PC-ko (Figure 2c). These data demonstrate that MPNSTs lacking plasma cells are no longer sensitized by CDK4/6-MEK inhibition to undergo extended tumor regression in response to anti-PD-L1 therapy.

The inability of ICB therapy to suppress MPNST growth in the absence of plasma cells directly correlated with reduced survival of those mice. Differences in the response to ICB therapy between wild-type and PC-ko mice were seen as early as day 10 post-therapy where only 69% of PC-ko mice treated with anti-PD-L1 alone were alive versus 100% of wild-type mice (Figure 2b, top). At day 28, no PC-ko mice receiving PD-L1 monotherapy were alive compared to 15% of wild-type mice (Figure 2b, bottom). Likewise, for treatment with CDK4/6-MEK inhibitors plus anti-PD-L1 therapy, only 22% of PC-ko mice were surviving at day 28 with no tumors regressing whereas 77% of wild-type mice were alive with 3 of 9 tumors still regressing (Figure 2b, bottom). Kaplan-Meier analysis similarly revealed that loss of plasma cells (bottom graph) abolished the extended survival and ∼10% cure rate obtained with anti-PD-L1 plus CDK4/6-MEK inhibition in wild-type mice ^20^ (Figure 2d). Moreover, while survival curves for anti-PD-L1 monotherapy and CDK4/6-MEK inhibitor therapy overlapped in wild-type mice, they were statistically different in PC-ko mice with the survival rate for anti-PD-L1 monotherapy now overlapping with vehicle plus IgG controls (Figure 2d).

Together, these data show that kinase inhibitor therapy concurrently targeting CDK4/6 and MEK is equally effective against *de novo* MPNSTs regardless of plasma cell status. By comparison, plasma cells are essential for an effective MPNST response to anti-PD-L1 ICB therapy, whether as a single agent or when combined with CDK4/6-MEK inhibition.

### Plasma cell-dependent differential gene expression at the single cell level induced by CDK4/6-MEK inhibition in *de novo* MPNSTs

Because MPNSTs in PC-ko mice were not sensitized to ICB therapy by CDK4/6-MEK inhibition, we reasoned that plasma cells must be required to establish a favorable immune environment that primes the tumor for successful immunotherapy. To test that idea, moderately sized *de novo* MPNSTs (∼450 mm^3^) in wild-type and PC-ko mice were treated for 4-6 days with CDK4/6 and MEK inhibitors. This allowed regressing tumors of sufficient size to be harvested for single cell RNA sequencing (scRNA seq) and molecular analyses (Figure 3a). *FindAllMarkers* ^42^ was used to identify cell populations from processed single cell transcriptomes from two wild-type tumors and three PC-ko tumors. The major cell populations identified within the tumors included MPNST cells, myeloid cells, T cells, B/Plasma cells, vascular endothelium, lymphatic endothelium, pericytes, and myocytes (Figure 3b). Select transcripts used to define each population are listed in Supplementary Figure 1.

**Figure 3.**
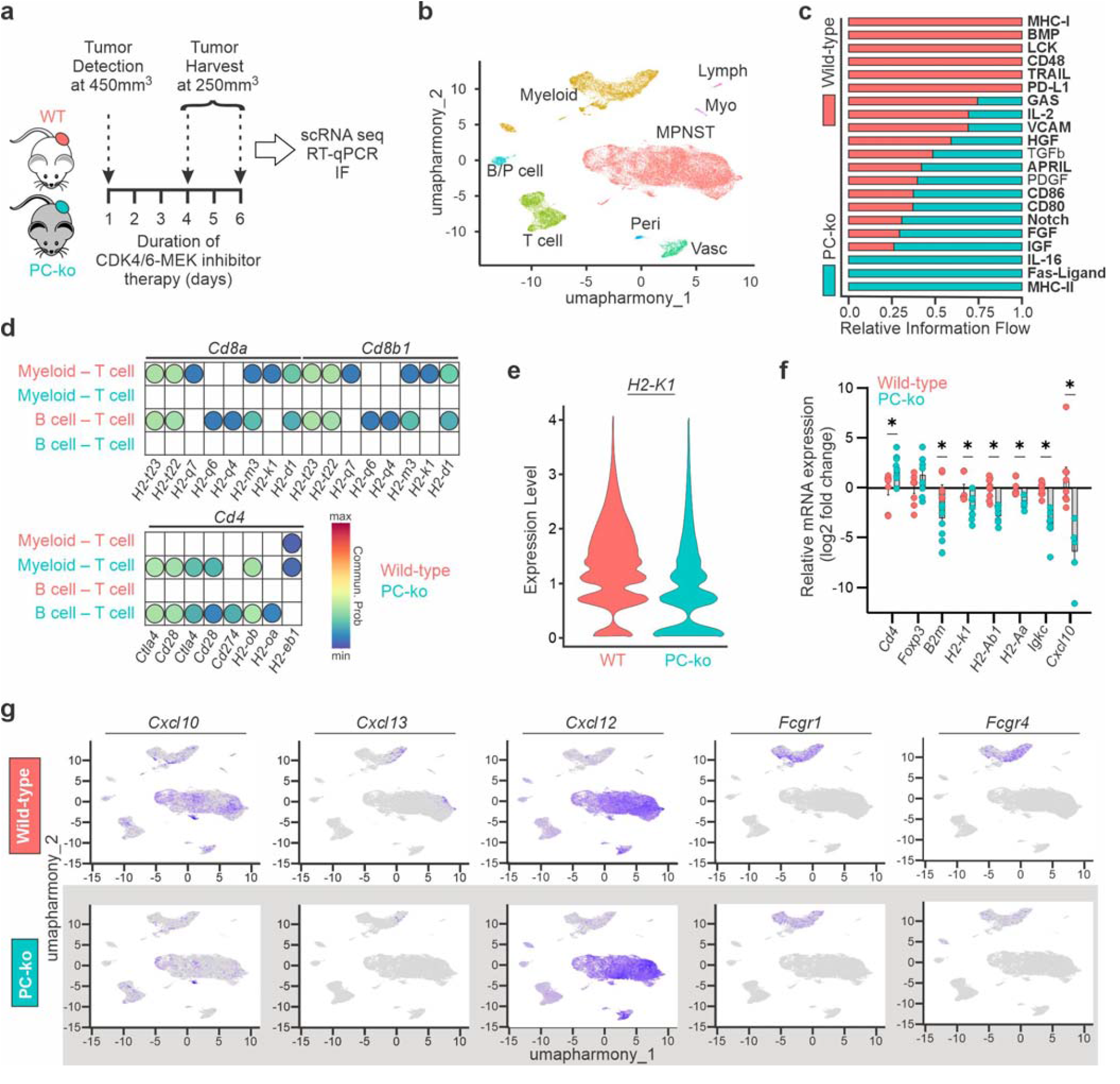
Loss of plasma cells reduces antigen presentation and predicted engagement of MHC-I by CD8. Wild-type (WT) and PC-ko MPNSTs, initiated by *Nf1/Ink4a/Arf* editing in the sciatic nerve, were treated daily with CDK4/6-MEK inhibitors (palbociclib at 100 mg/kg, mirdametinib at 1 mg/kg) once tumors reached 450 mm3. Tumors were harvested between days 4 to 6 of therapy when they regressed to 250 mm3. a) Schematic of experimental design. b) Visualization of identified cell types from scRNA seq. c) Top predictions from CellChat for differentially regulated receptor-ligand interactions in WT and PC-ko tumors. d) Focused bubble plots for CellChat predicted interactions between the indicated ligands and receptors expressed on the Myeloid/B cells and T cells. Commun. Prob., communication probability. e) Relative expression levels of H2-k1 within the MPNST cluster in WT and PC-ko MPNSTs. f) Quantitative RT-qPCR validation of selected genes suggested by scRNA seq to be up in WT or PC-ko tumors. Relative mRNA expression was calculated using the delta delta CT method. g) Feature plots for visualization of the expression levels of selected transcripts overlayed onto panel b. Error bars are SEM. P value was obtained via Student’s t test (*, P < 0.05)

We first focused on plasma cell-dependent gene expression intrinsic to the MPNST population. Non-negative matrix factorization (NMF) was employed to identify metaprograms (MPs), groups of co-differentially expressed genes (DEGs) (Supplementary Figure 2a), which represent recurrent programs of intra-tumoral heterogeneity (Supplementary Figure 2b). Eight MPs were defined, and the top 50 genes in each were extracted for MP identification and downstream analysis. Those metaprograms showing the greatest differences between wild-type and PC-ko MPNSTs were MP1, MP4, and MP5 (Supplementary Figure 2c). Wild-type tumors containing plasma cells expressed higher levels of transcripts within pathways related to embryonic/neuronal development (MP1) and B/Plasma cell immunity (MP4) whereas PC-ko tumors were enriched for pathways related to myeloid immunity/motility (MP5) (Supplementary Figure 2, d-f).

### Plasma cells promote MHC-I antigen presentation to CD8+ T cells and pro-inflammatory chemokine expression

The two most differentially expressed metaprograms were centered on professional antigen presenting cells, B cells (MP4) and myeloid cells (MP5), suggesting antigen presentation is regulated by plasma cells in MPNSTs (Supplementary Figure 2, b, c, e, and f). We used CellChat, a tool to infer and quantify intercellular receptor-ligand interactions from scRNA seq data, to evaluate interactions between major histocompatibility complex class I (MHC-I) or II (MHC-II) antigen presentation molecules and T cell co-receptors, CD8 and CD4 ^43^. Strong MHC-I and PD-L1 binding to CD8 and PD-1, respectively, on T cells was exclusive to wild-type tumors (Figure 3c). The opposite, MHC-II interactions with CD4+ T cells, defined plasma cell-deficient MPNSTs ^44^. Downstream analysis through CellChat infers the strength and cell types involved for each receptor-ligand interaction. MPNSTs in wild-type mice were enriched for interactions between many MHC-I molecules (*H2-t23, H2-k1*, etc.) on myeloid and B cells with *Cd8a/Cd8b1* T cells (Figure 3d, top). These interactions were absent in PC-ko MPNSTs, which instead presented with more interactions between *Cd4* T cells and MHC-II molecules (*H2-ob, H2-oa, H2-ab1*) (Figure 3d, bottom).

The predicted receptor-ligand interactions reflected differential expression of the antigen presentation and receptor mRNAs. This included elevated *H2-k1* MHC-I transcripts in wild-type tumors and increased *Cd4* in PC-ko tumors by single cell sequencing (Figure 3e) and RT-qPCR validation (Figure 3f), respectively. Unexpectedly, RT-qPCR revealed that PC-ko tumors not only have lower expression of MHC-I (*B2m, H2-k1*) transcripts but also reduced MHC-II molecules (*H2-ab1* and *H2-aa*) (Figure 3f). This suggests that plasma cells support antigen presentation through both MHC-I and MHC-II molecules, implying that the increase in predicted MHC-II-dependent interactions in PC-ko mice may be a product of increased CD4+ T cells.

Cytokines and chemokines shape the tumor microenvironment by directing immune cell infiltration into tumors. Dimensional reduction and visualization of the single cell sequencing data revealed elevated expression of *Cxcl10* and *Cxcl13* mRNAs in the MPNST and myeloid cells of wild-type tumors versus higher *Cxcl12* in MPNST and vascular cells of PC-ko tumors (Figure 3g). Both CXCL10 and CXCL13 chemokines contribute to a “hot” tumor microenvironment by recruiting and activating T and B cells, and their expression is required for and positively correlates with better responses to ICB therapy ^45-51^. By comparison, while CXCL12 can also attract immune cells as part of an inflammatory response, its binding to the CXCR4 receptor activates signaling that promotes tumor growth, angiogenesis, metastasis, and survival ^52^. Consistent with the pro-inflammatory landscape of wild-type tumors, transcripts for *Fcgr1* and *Fcgr4* (low affinity receptors for IgG proteins), were upregulated in myeloid cells of wild-type MPNSTs. Both Fcγ receptors are associated with increased tumor infiltration and activation of immune cells including T cells, and engage with anti-PD-L1 antibodies to augment their *in vivo* activity ^53,54^

### Plasma cells promote the infiltration and activation of CD8+ T cells within CDK4/6-MEK inhibited MPNSTs

Immune checkpoint blockades, such as those targeting PD-L1 and PD-1, work by sustaining the activity of anti-tumor T cells ^55^. To better understand how plasma cells promote T cell activation, we isolated the T cell cluster from our scRNA seq dataset for further analysis. Dimensional reduction and visualization identified multiple transcriptionally distinct subpopulations of T cells with unique patterns of enrichment in wild-type and PC-ko tumors (Figure 4a). Following clustering on this subpopulation of T cells and identification of DEGs, we identified seven major subpopulations of lymphoid cells which included CD4+ T cells, CD8+ T cells, natural killer (NK) cells, gamma delta T (γδT) cells, NKT cells, regulatory T (Treg) cells, and Interferon-induced T cells (IFI T) (Figure 4b). Select genes used to delineate subtypes are shown in Supplementary Figure 3a.

**Figure 4.**
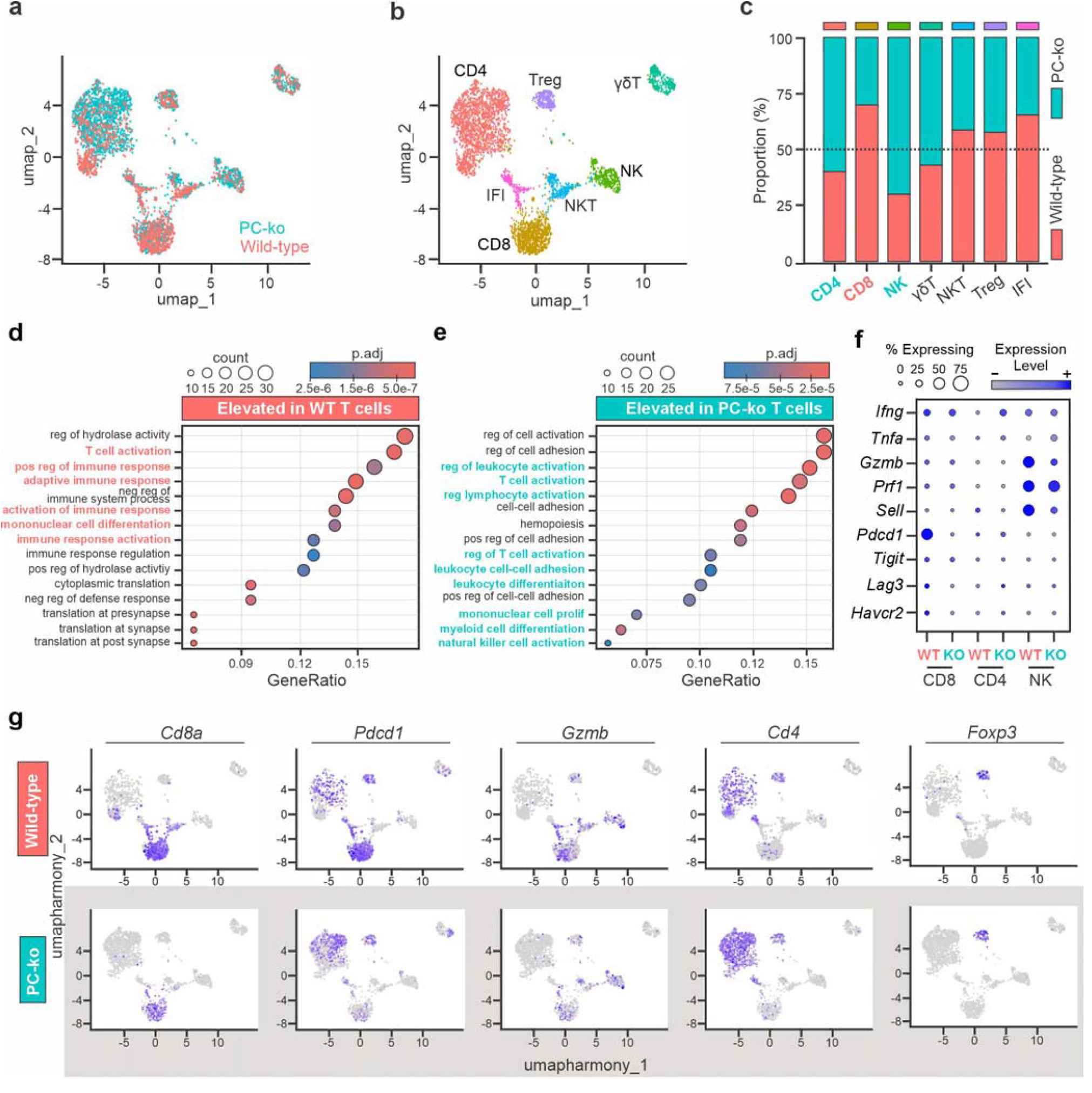
Plasma cells support the activation of CD8+ T cells. Analysis of the T cell cluster identified by FindAllMarkers in scRNA seq data from wild-type and PC-ko MPNSTs that were generated and treated as shown in Figure 3a. a) Dimensional reduction and visualization of the T cell cluster split by genotype. b) FindAllMarkers was used to identify T cell subtypes. c) Relative amounts of T cell subtypes quantified in WT versus PC-ko mice. Gene Set Enrichment Analysis of biological pathways upregulated in T cells from d) WT or e) PC-ko tumors. f) Relative expression levels of selected T cell exhaustion and activation markers in CD8+ T cells, CD4+ T cells, and NK cells. g) Feature plots for visualization of selected transcripts overlayed onto panels a and b.

In alignment with CellChat predictions, significantly more *Cd8*-expressing T cells were present in wild-type tumors compared to PC-ko tumors (Figure 4c). This was mirrored by increases in *Cd4*-expressing T cells as well as NK cells within PC-ko tumors (Figure 4c). Gene ontology (GO) analysis revealed that T cells from wild-type tumors were enriched for biological pathways related to T cell activation and the adaptive immune response (Figure 4d). PC-ko tumors also displayed enrichment of T cell activation pathways, as well as gene signatures of activated NK and myeloid cells (Figure 4e).

CD8+ T cells are known to be activated in response to CDK4/6 and MEK inhibition ^26^. However, an enrichment of T cell activation gene signatures in both wild-type and PC-ko T cells could indicate that plasma cells mediate the activation of specific T cell subsets in response to combined CDK4/6 and MEK inhibition. To assess that possibility, we evaluated the expression of transcripts associated with T cell activation within the T cell cluster. Consistent with cell chat predictions of CD8+ T cell activation, a large fraction of CD8+ T cells from wild-type tumors expressed high levels of *Pdcd1*, which encodes the checkpoint protein, programmed cell death protein 1 (PD-1) (Figure 4, f and g). PD-1 can be a marker of both T cell activation and exhaustion ^56^, therefore we examined the expression of other classical immune checkpoint proteins, Lag3 (*Havcr2)*, Tim3, and Tigit. These exhaustion marker mRNAs were expressed at very low levels throughout all T cell subsets, indicating minimal T cell exhaustion in both wild-type and PC-ko tumors at the time of analysis (Figure 4f). Interestingly, while NK cell numbers were elevated in PC-ko tumors, NK cells in wild-type tumors expressed higher levels of Granzyme B (*Gzmb)* and CD62L (*Sell*). Those changes suggest greater NK cell activation and maturation ^57,58^ in wild-type tumors.

The expression of select T cell transcripts was visualized via feature plots of the T cell population. In agreement with the above findings, increased *Cd8a* expression marked a larger population of CD8+ T cells in wild-type tumors whereas increased *Cd4* defined an enriched CD4+ T cell population in PC-ko tumors (Figure 4g). *Pdcd1* expression was primarily elevated in CD8+ T cells within wild-type tumors, which also expressed increased *Gzmb* (component of cytotoxic granules that marks activated T [and NK] cells). Some CD4+ T cells expressed *Pdcd1* in PC-ko tumors, but at markedly lower levels than in CD8+ T cells from wild-type tumors. *Foxp3*, a marker of regulatory CD4+ T cells was unchanged between wild-type and PC-ko tumors. These results suggest that plasma cells are essential for the specific recruitment and activation of CD8+ T cells in CDK4/6-MEK inhibitor treated MPNSTs.

### Plasma cells support antigen presentation in myeloid cells within CDK4/6-MEK inhibited MPNSTs

The presentation of antigen, either at the tumor cell surface through MHC-I or through antigen presenting cells (APCs) expressing MHC-II, is a critical, early step in T cell activation. We isolated the two main APC clusters from our scRNA seq dataset for characterization: B/plasma cells and myeloid cells. On the B/plasma cell cluster (Figure 5A), unbiased clustering delineated B cells from plasma cells and confirmed that plasma cells are only present in wild-type tumors (Figure 5b). Gene sets used to determine B and myeloid cell subtypes are listed in Supplementary Figure 3, b and c, respectively.

**Figure 5.**
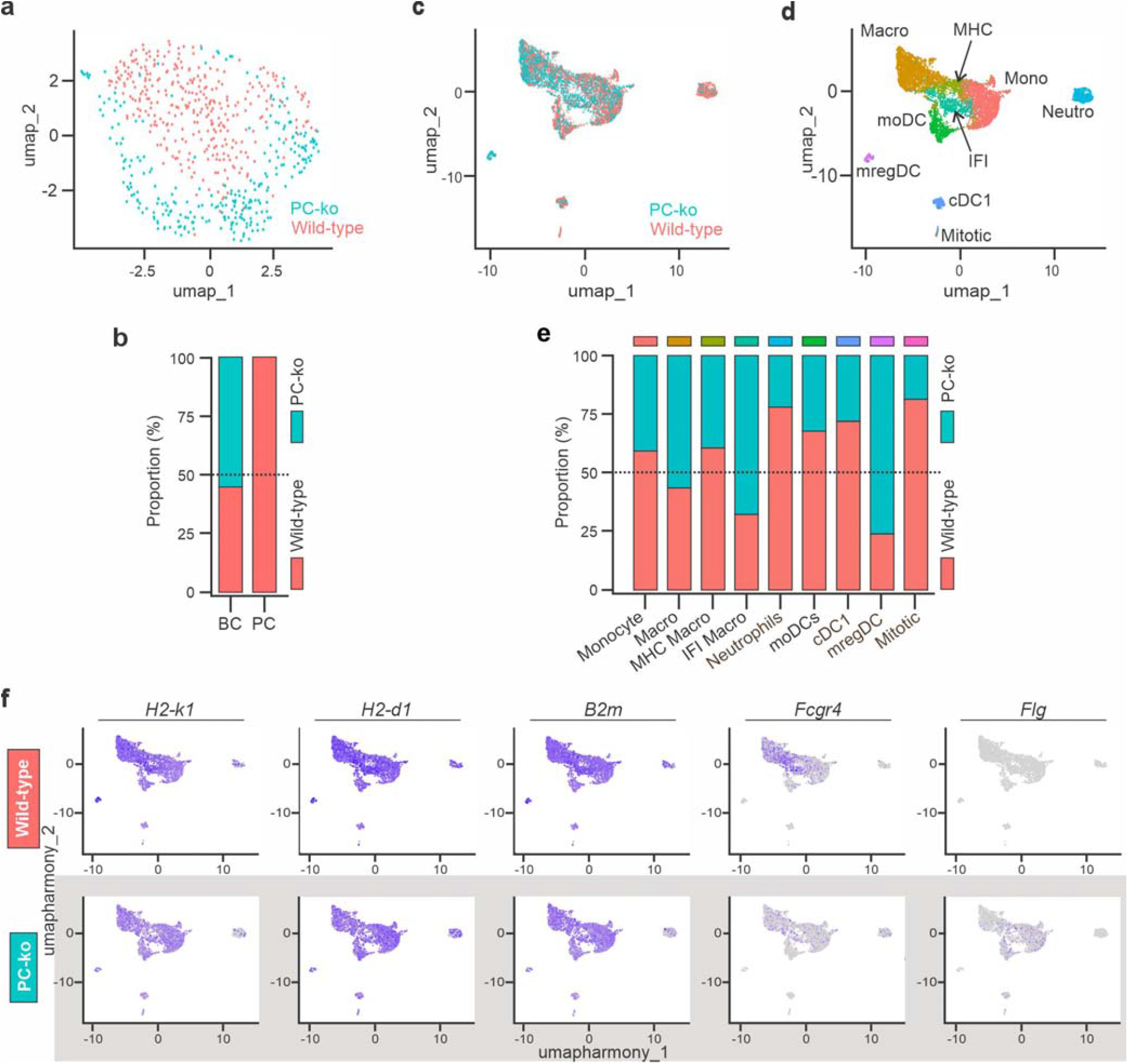
Plasma cells support antigen presentation and expression of antibody receptors on myeloid cells in CDK4/6-MEK inhibited *de novo* MPNSTs. B/Plasma cell and myeloid clusters were isolated from scRNA seq data for further analysis. a) Dimensional reduction and visualization of B/Plasma cell clusters. b) Identified subsets are listed and relative amounts in wild-type and PC-ko MPNSTs are quantified. c) Dimensional reduction and visualization of the Myeloid cluster. Identified cell types are visualized in d) with relative amounts of each subtype quantified in wild-type versus PC-ko MPNSTs shown in e). f) UMAP Feature plots for visualization of select transcript expression overlayed onto the myeloid population (see panel c).

Myeloid cells play important roles in MPNST biology and comprise a large portion of the tumor microenvironment ^59-62^. The myeloid compartment of MPNSTs is complex and comprised of many cell subpopulations (Figure 5, c and d). Following clustering and DEG identification, nine main subtypes of myeloid cells were defined in our scRNA seq dataset: monocytes, macrophages (Macro), MHC high macrophages (MHC Macro), Interferon-induced macrophages (IFI Macro), Neutrophils, monocyte-derived dendritic cells (moDCs), classical dendritic cells (cDC1), regulatory dendritic cells (mregDC), and mitotic myeloid cells (Figure 5d).

Many cell types were not affected by plasma cell status, including monocytes, general macrophages, and MHC-high macrophages. Among the altered myeloid populations, neutrophils, moDCs, and cDC1s were significantly reduced in PC-ko tumors (Figure 5e), suggesting that plasma cells support their presence in tumors. Conversely, PC-ko tumors had enriched mregDCs (Figure 5e), which famously express immunosuppressive cytokines that promote resistance to ICB therapy ^63,64^. Visualization of MHC-I molecules *H2-k1, H2-d1*, and *B2m* across all myeloid subsets showed reduced expression of all three transcripts in PC-ko tumors (Figure 5f). Additionally, myeloid cells from wild-type tumors expressed more *Fcgr4*, which can aid in the uptake of immune complexes by binding the constant region of the antibody. In contrast, myeloid cells in PC-ko MPNST expressed modest levels of *Flg*, which was undetectable in myeloid cells within wild-type tumors. *Flg* encodes filaggrin, a keratin filament binding protein that provides structural support to the skin and maintains its barrier functions ^65^. Of note, recent reports in urothelial carcinoma show that *Flg* expression predicts poor patient response to anti-PD-1 therapy ^66^. Together, these data indicate that plasma cells remodel the myeloid compartment to promote antigen presentation and a pro-inflammatory microenvironment.

### Histological validation of intratumoral immune cell abundance in CDK4/6-MEK inhibited MPNSTs

Single cell RNA seq findings were validated at the cellular level and protein levels using immunofluorescent staining of immune cell subsets in the same wild-type and PC-ko MPNSTs that were treated with CDK4/6-MEK inhibitors for 4 to 6 days. As expected, only wild-type MPNSTs contained kappa-light chain positive (KLC+) cells (Figure 6a), a specific marker for plasma cells ^67^. In alignment with the scRNA seq data, wild-type MPNSTs had more intratumoral CD8+ T cells (nearly 3-fold) whereas PC-ko tumors had more CD4+ T cells (Figure 6a). Quantification of positive cells verified statistically significant differences for each cell type in wild-type versus PC-ko tumors (Figure 6b).

**Figure 6.**
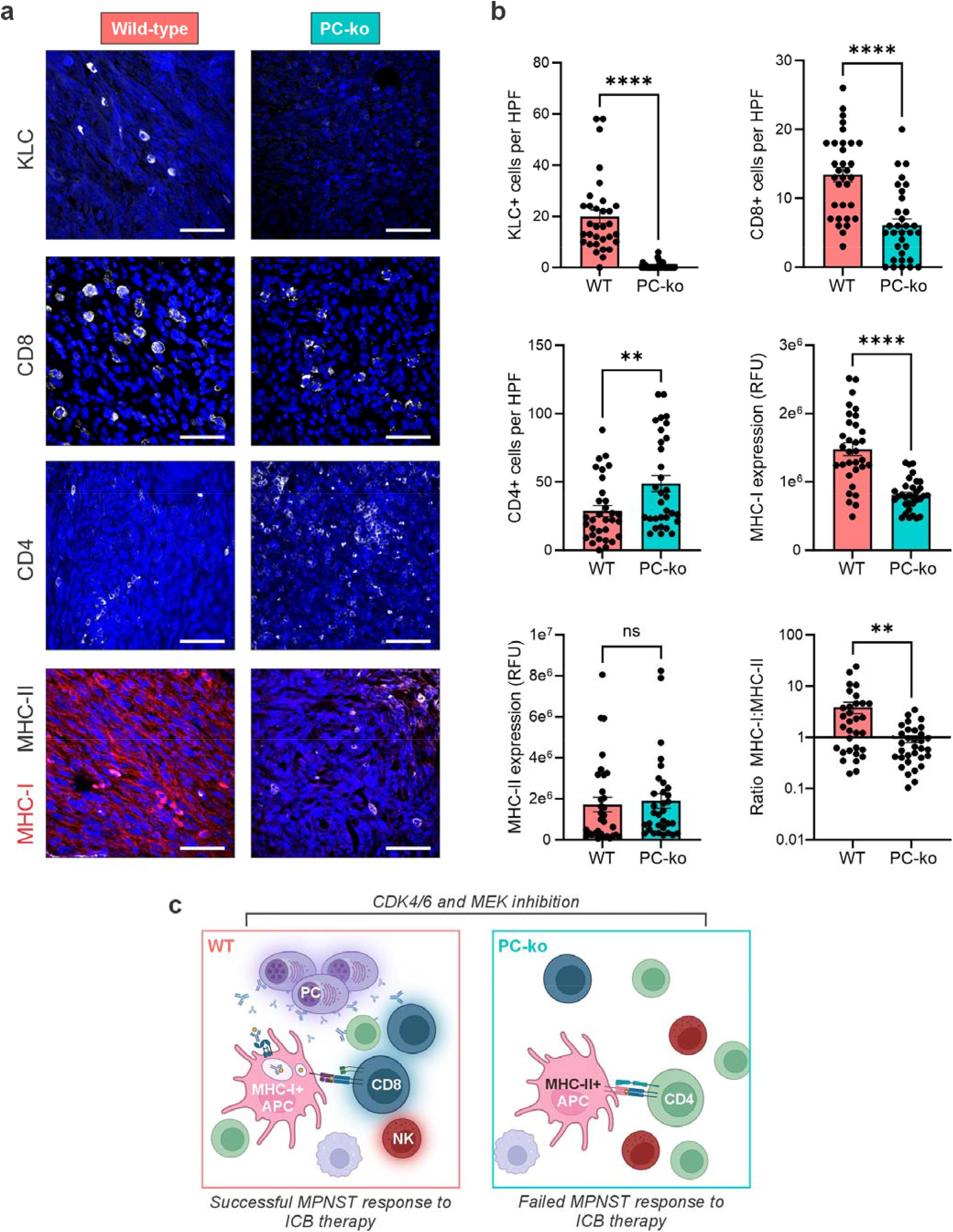
Plasma cells support MHC-I expression and intratumoral CD8+ T cell populations. Wild-type and PC-ko MPNSTs were generated and treated as illustrated in Figure 3a. a) Representative 40X images showing immunofluorescence (IF) staining of Kappa Light Chain (KLC), CD8, CD4, MHC-I, and MHC-II in de novo MPNSTs. White scale bar represents 50 µM. b) Quantification of IF results from A. Error bars are SEM. P value was obtained via Student’s t test (ns, non-significant; **, P < 0.01, ****, P < 0,0001). c) Model depicting distinct immune environments in plasma cell positive (WT) versus negative (PC-ko) MPNSTs following CDK4/6-MEK inhibition. Successful ICB therapy in WT MPNSTs requires plasma cells (PCs) that direct antigen presenting cells (APCs) to display antigen to CD8+ T cells through MHC-I. PC-ko MPNSTs, which fail to respond to ICB therapy, are characterized by an MHC-II+ APC interaction with CD4+ T cells and an insufficient CD8+ response. The glow surrounding cells indicates activation.

Similarly, as predicted from the single cell sequencing data, significantly higher expression of MHC-I protein was seen in wild-type MPNSTs (Figure 6, a and b). MHC-II protein was not statistically different between the tumor types although it trended to higher levels of expression in PC-ko tumors (Figure 6, a and b). As a result of these differences, wild-type tumors exhibited an increased ratio of MHC-I to MHC-II protein expression relative to PC-ko MPNSTs (Figure 6b). H&E stained sections of the tumors were analyzed for potential differences in lymphoid aggregates at the histological level. Immune cell aggregates appeared as diffuse clusters in both wild-type and PC-ko tumors with an overall absence of highly organized regions such as germinal centers or tertiary lymphoid structures (Supplementary Figure 4a). The total lymphoid aggregate area was quantified and found to be similar in both wild-type and PC-ko MPNSTs (Supplementary Figure 4a). These findings reveal that plasma cells dictate the composition of the inflammatory immune landscape in CDK4/6-MEK targeted MPNSTs (CD8+ T cells with MHC-I in wild-type versus CD4+ T cells in PC-ko).

It is well known that the MPNST microenvironment is dominated by tumor associated macrophages (TAMs) ^62^. No changes were seen in the total number of CD68+ tumor associated macrophages (TAMs) between wild-type and PC-ko tumors (Supplementary Figure 4b, top), indicating that plasma cells are not vital for sustaining a TAM population. Interestingly, wild-type MPNSTs displayed slightly higher PD-L1 expression compared to PC-ko MPNSTs (Supplementary Figure 4b, bottom), which supports the predicted increase in PD-1/PD-L1 signaling in wild-type tumors by CellChat analysis (Figure 3c). Representative images of negative controls for each immunofluorescent stain are depicted in Supplementary Figure 4c.

As modeled in Figure 6c, the single cell transcriptome and cell-based IF data support the conclusion that plasma cells direct antigen presentation to CD8+ T cells through MHC-I in MPNSTs, which corresponds with successful response to ICB therapy. In the absence of plasma cells, however, tumor cell antigens are primarily presented through MHC-II to CD4+ T cells and there is a failed response to ICB therapy.

### B and plasma cell abundance prognosticates enhanced survival in human sarcomas

To assess the importance of plasma cells in sarcoma patient outcome, we queried The Cancer Genome Atlas Sarcoma (TCGA SARC) bulk RNA sequencing (RNA-seq) dataset and predicted relative immune cell subsets using the CIBERSORT algorithm ^68^. The cell fraction for each subtype was identified for all sarcomas while samples without a detectable fraction of plasma cells were excluded from downstream analysis. In many other cancers, plasma cells co-organize with immune cells that promote anti-tumor immunity including B cells, CD4+ helper T cells, and CD8+ cytotoxic T cells ^28,29^. In agreement with this, the plasma cell fraction in sarcomas positively correlated with each of those cells while being inversely correlated with pro-tumorigenic cell types, such as M2 macrophages and mast cells (Figure 7a).

**Figure 7.**
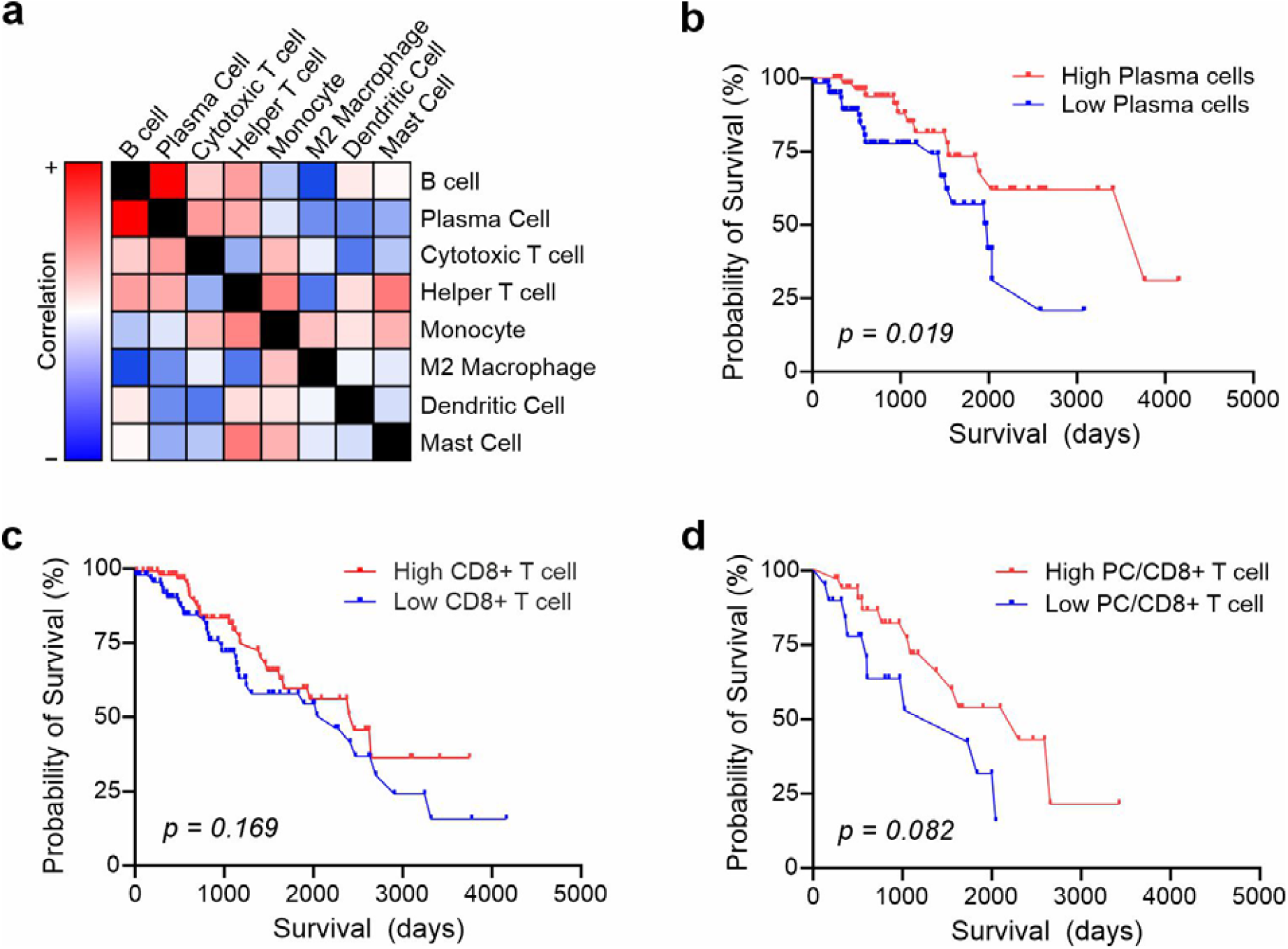
Plasma cells correlate with CD8+ T cells in patient sarcomas and predict prolonged patient survival. CIBERSORT was used to deconvolute and predict immune cell subsets from publicly available RNA sequencing data in TCGA SARC. a) Pearson correlation matrix of immune cell subsets with each other. Red, direct correlations (r = 1); blue, indirect correlation (r = −1). Panels b-d: Kaplan Meier survival analyses based on gene signatures for b) plasma cells, c) T cells, and d) both plasma cells and T cells. Any samples with null values for either immune cell signature were removed and remaining samples were bisected at the upper and lower 45th percentile. P values are indicated.

Intratumoral plasma cells are associated with prolonged survival in most cancers ^27-30^, prompting us to perform a Kaplan-Meier survival analysis of the TCGA SARC dataset. Samples were binned into the upper and lower 45^th^ percentiles based off their relative plasma cell fraction. Sarcoma patient survival was significantly extended in patients with a higher intratumoral plasma cell signature (Figure 7b). Surprisingly, a higher fraction of CD8+ T cells did not correlate with longer patient survival (Figure 7c). Because CD8+ T cells and plasma cells are proportional with each other in these tumors (Figure 7a), we then compared the survival of patients with high versus low fractions of both cell types. Patients with elevated intratumoral CD8+ T cell and plasma cell signatures trended to having longer survival times than those with lower fractions, although this difference did not reach statistical significance (Figure 7d). These findings suggest that increased intratumoral plasma cells are more valuable prognostic markers than CD8+ T cells for prolonged patient survival in sarcomas.

## Discussion

MPNSTs have an immune suppressive tumor microenvironment and do not respond well to any therapy, including ICB immunotherapy. We recently reported that MPNSTs can be sensitized to anti-PD-L1 therapy by CDK4/6-MEK inhibition, which uniquely stimulates plasma cell accumulation in the tumors coincident with CD8+ T cell activation. Here, we directly tested the importance of plasma cells to the ICB response in MPNSTs and found they are vital for successful immunotherapy.

This is the first study in any cancer type to demonstrate a causal relationship between plasma cells and the tumor immune response to ICB therapy. In many cancers, B and plasma cells are increasingly recognized as robust prognosticators of prolonged patient survival ^27-30^ and improved response to ICB agents ^28,31-34^. However, the role of plasma cells in therapy-induced anti-tumor immunity has never been investigated and only a few studies have explored the contributions of B cells. Genetic ablation or antibody depletion of all B-lineage cells greatly reduced the anti-tumor effects of ICB monotherapy as well as ICB combination therapies in preclinical tumor models ^21,39^. Griss *et al*. evaluated melanoma patients and likewise found that B cells predicted response to ICB therapy and were required for establishing a pro-inflammatory state in the tumors ^40^. They found that newly differentiated B cells, called plasmablasts (which are lost in B cell deficient mice), were crucial for producing cytokines for T cell recruitment. Plasmablasts not only secrete cytokines, they also make antibodies (albeit at lower levels than mature plasma cells) and internalize antigens for presentation to T cells (a function lost by mature plasma cells) ^69,70^. Because plasmablasts are still present in *AID−/−; µS−/−* mice, which only lack mature plasma cells, our data suggest it is the mature plasma cells that mediate the anti-tumor response to anti-PD-L1 therapy in MPNSTs.

Plasma cells are best known for producing antibodies, which carry out many potential roles in the tumor microenvironment (TME) that are incompletely understood ^71^. First, antibodies can activate the complement cascade, which causes immunologic, osmotic death through the formation of pores in the cell membrane ^72^. Second, antibodies can promote lysis of target cells by triggering natural killer (NK) cell activation. That process, termed antibody-dependent cellular cytotoxicity (ADCC), relies on the binding of antibody Fc regions to Fcγ receptors on NK cells ^73^. Indeed, *Fcgr4* was upregulated in wild-type MPNSTs and encodes FcgRIV, an essential mediator of ADCC ^74^. Finally, antibodies can label tumor cells for phagocytosis or form immune complexes to aid antigen presentation by phagocytes ^72^. Many of these pathways were selectively upregulated in wild-type, CDK4/6-MEK targeted MPNSTs according to NMF analysis, but not in PC-ko tumors. Thus, intratumoral plasma cells may support CD8+ T cell activation by forming immune complexes with antigen for uptake and presentation by APCs. In addition to antibody-dependent mechanisms in which plasma cells remodel the TME, they may secrete other pro-inflammatory factors to promote immune cell infiltration, increase MHC-I expression, and enhance anti-tumor immunity.

As MPNSTs develop, they become increasingly “cold” immunologically with reduced antigen presentation, exclusion and/or exhaustion of infiltrating T cells, and preponderance of pro-tumorigenic M2 macrophages, which promotes tumor resistance to ICB therapies ^75^. We found that dual CDK4/6-MEK inhibition for just 4 to 6 days in wild-type MPNSTs was sufficient to activate a pro-inflammatory, MHC-I directed CD8+ T cell response. That rapid remodeling of the immune milieu favorably corresponded with sustained anti-tumor effects of anti-PD-L1 therapy and significantly improved survival. Conversely, MPNSTs lacking plasma cells had an MHC-II, CD4+ T cell phenotype and were unresponsive to anti-PD-L1 therapy. An enriched CD8+ T cell response and bolstered antigen presentation have been noted in other tumor types sensitized to immune checkpoint blockades, either through chemotherapy and MEK inhibition ^76^ or epigenetic therapies ^77^. Others have also shown that combined inhibition of CDK4/6 and MEK triggers a strong anti-tumor response in Ras-driven pancreatic adenocarcinoma that depended on CD8+ T cell activation ^26^. Plasma cells were not examined in the above models, but our work suggests that they likely mediated or contributed to the successful remodeling of the immune landscape and sensitization to ICB therapies in those settings.

Transcripts expressing chemokines (*Cxcl10, Cxcl13*) and antigen presentation molecules (*H2-k1, B2m, H2-Aa*) were notably upregulated in wild-type, plasma cell-containing MPNSTs treated with CDK4/6 and MEK inhibitors relative to tumors lacking plasma cells. Those intratumoral changes are widely known to support infiltration and activation of immune cells (B, T, myeloid, and dendritic cells) which may, over time, assemble with plasma cells into mature tertiary lymphoid structures (TLS) that heighten CD8+ T cell activity. Histological analyses of the CDK4/6-MEK targeted MPNSTs revealed loosely clustered lymphoid aggregates of similar morphology and size in both wild-type and plasma cell-deficient mice. We observed no evidence of bona fide TLS formation. We speculate the diffuse lymphoid aggregates detected are pre-TLS in nature, suggesting that 4 to 6 days of kinase inhibitor therapy is an insufficient length of time to induce TLS formation. It will be interesting to determine in future studies if the number and/or location of intratumoral plasma cells within kinase inhibitor-induced TLSs (emerging versus mature) dictates the magnitude and duration of successful ICB therapy.

Histologic detection of tumor infiltrating immune cells and CellChat intercellular receptor-ligand interaction analyses indicate that CD8+ T cell infiltration and activation in CDK4/6-MEK targeted MPNSTs is a plasma cell dependent process. Two of the strongest receptor-ligand interactions observed exclusively in wild-type tumors were for CD8+ T cell receptor associations with MHC-I and lymphocyte-specific protein tyrosine kinase (LCK). LCK plays a central role in T cell activation and anti-tumor immunity. It is critical for remodeling the TME in melanoma and predicts response to immunotherapy ^78^. Across multiple cancers, LCK levels positively correlate with CD8+ T cell tumor infiltration ^79^. Tumor necrosis factor-related apoptosis-inducing ligand (TRAIL), which had strong predicted interaction with death receptors in wild-type MPNSTs, is upregulated on activated T cells and contributes to T cell killing of tumor cells ^80^. Conversely, dominant receptor-ligand interactions in PC-ko MPNSTs included MHC-II with CD4, IL-16 (a chemoattractant for CD4+ cells), and Fas ligand (FasL), which is highly expressed in some MPNSTs and has a complex role in anti-tumor immunity. Endothelial FasL is linked with an absence of CD8+ T cells due to preferential killing of those cells ^81^. Defining the essential mechanisms by which plasma cells augment the T cell response may be crucial for delineating which MPNSTs or other solid tumors are likely to respond well to ICB therapy.

In addition to the immunological effects of plasma cells in MPNST therapy response, we saw increased embryonic and neuronal development gene expression in wild-type tumors. This could reflect an unwanted increase in stem-like tumor cells within the drug-treated MPNSTs. By activating CD8+ T cells and effectively clearing a greater portion of the tumor cells, CDK4/6-MEK inhibition could cause an undesirable enrichment for a stem-population less affected by cell cycle-targeting therapies, potentially driving drug resistance. Other transcripts associated with cancer stem cells, like *Fos* ^82^ and *Dusp1* ^83^, were concurrently elevated. Our data also predict an upregulation of hepatocyte growth factor (HGF) signaling in wild-type tumors. HGF signals through its receptor, MET, a tyrosine kinase receptor which is a marker of stemness in glioblastoma ^84^, is genetically amplified along with *HGF* in NF1-related MPNSTs ^85^, and is sufficient for Schwann cell dedifferentiation into MPNSTs ^14^. These observations are relevant because most tumors in wild-type mice (85-90%) treated with CDK4/6, MEK, and PD-L1 targeted therapy ultimately develop resistance and regrow even though 10-15% of mice are cured (^20^ and data herein). It is possible that therapy-resistant stem-like MPNST cells residing in the TME are enriched upon T cell destruction of other tumor cells and then reinvigorate the MPNST population. Further analysis of therapy sensitive versus resistant MPNSTs should provide valuable insights into the role of stem-like MPNST cells in resistance to therapies targeting CDK4/6, MEK, and PD-L1.

In summary, we show that plasma cells are required for MPNSTs to respond to anti-PD-L1 therapy. Tumor resident plasma cells function, at least in part, by remodeling the MPNST immune landscape into a pro-inflammatory, CD8+ T cell milieu with increased MHC-I antigen presentation. That immune state is primed for an efficacious response to ICB agents. Our findings highlight two key points. First, tumor resident plasma cells may be pivotal in predicting which patients will respond to ICB therapies. Second, anticancer treatments that increase intratumoral plasma cells may be a key step in converting cold tumors into “hot” tumors that will respond effectively to ICB therapy. These points have increasing clinical relevance that likely extends beyond MPNSTs to other solid tumors because trials involving ICB therapies, particularly combinations with chemotherapeutic or targeted agents, are on the rise.

## Author Contributions

JJL and DEQ conceived and designed the studies. JJL, CA, JAR, CAK, ECE, and EV conducted experiments and acquired data. JJL, RR, JAR, EV, DKM, PB, RDD, JCH, BWD, and DEQ analyzed the data. RR, AJ, CRW, AWB, and NJK provided key reagents. JJL, RDD, BWD, and DEQ obtained funding. JJL, RR, DKM, RDD, JCH, BWD, and DEQ provided observations and scientific interpretations. All authors reviewed and approved the manuscript.

## Acknowledgements

We thank Dr. Kevin Knudtson and members of the Genomics Division of the Iowa Institute of Human Genetics at the University of Iowa Carver College of Medicine and Holden Comprehensive Cancer Center for scRNA sequencing. We also thank Drs. Snehajyoti Chatterjee (University of Iowa), Budhaditya Basu (University of Iowa), and Sara Gosline (Pacific Northwest National Laboratory) for providing insights on analysis of scRNA seq datasets.

## Data Sharing

Mouse tumor single cell RNA seq data will become publicly available in the Gene Expression Omnibus (GSE306038) upon publication. All other data generated in this study are available upon request from the corresponding author.

