## Supplementary Data for "Intratumoral plasma cells mediate CD8+ T cell infiltration and successful immune checkpoint blockade therapy in *de novo* MPNSTs"

Supplementary Figure 1

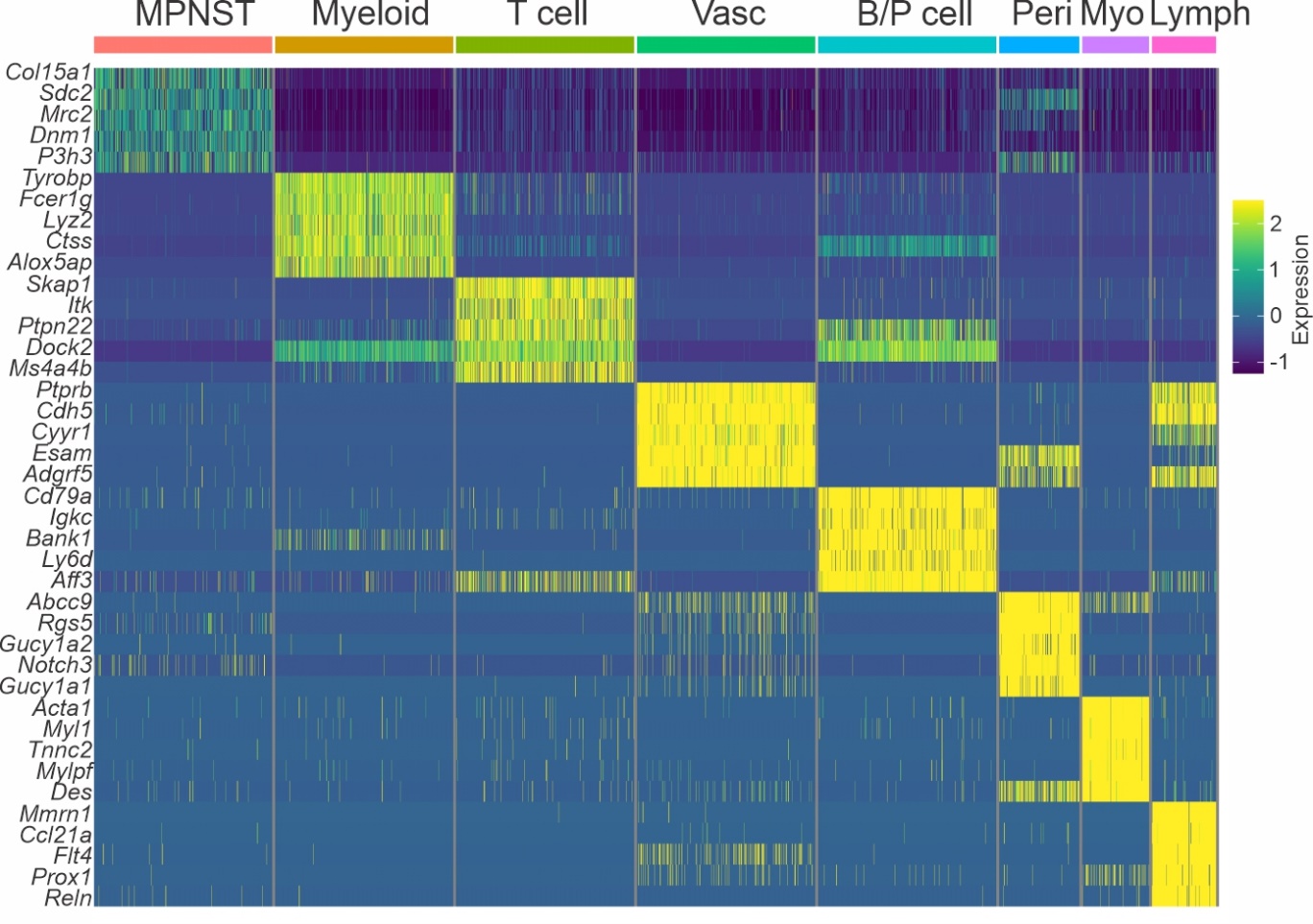

**Supplementary Figure 1. Identification and delineation of major cell populations in *de novo* MPNSTs.** Heatmap showing gene expression subsets, obtained from FindAllMarkers analysis of scRNA seq data, which were used to define the different major cell populations in *de novo* MPNSTs from WT and PC-ko mice that had been treated for 4 to 6 days with CDK4/6 and MEK inhibitors.

Supplementary Figure 2

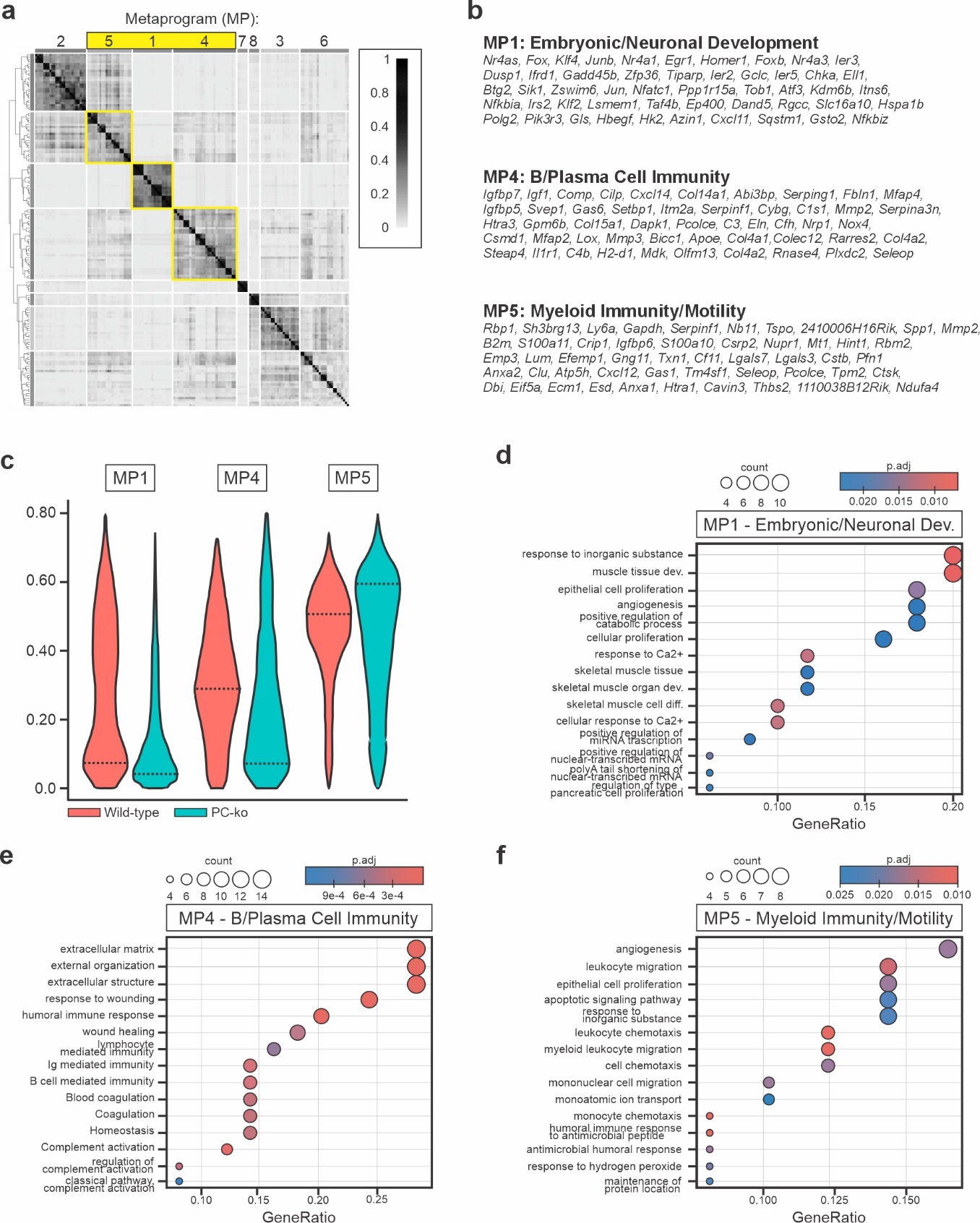

**Supplementary Figure 2.** **Non-negative matrix factorization of MPNST population clusters.** Non-negative matrix factorization (NMF) of scRNA seq data was employed to identify co-occurring panels of transcripts in WT and PC-ko MPNSTs that had been treated for 4-6 days with CDK4/6-MEK inhibitors. **a)** Greyscale heatmap of identified metaprograms (MPs) by NMF analysis with yellow highlighting those (MP1, MP4, and MP5) with significant differences between WT and PC-ko tumors. **b)** List of the top 50 genes from MPs 1, 4, and 5. **c)** Relative expression levels of MP1, MP4, and MP5. **Panels d-f:** Dot plots showing enriched pathways based off the expression of the transcripts for MPs 1, 4 and 5, as determined by gene set enrichment analysis (GSEA). **d)** MP1 – Embryonic/Neuronal Dev (development). **e)** MP4 – B/Plasma Cell Immunity. **f)** MP5 – Myeloid Immunity/Motility.

Supplementary Figure 3

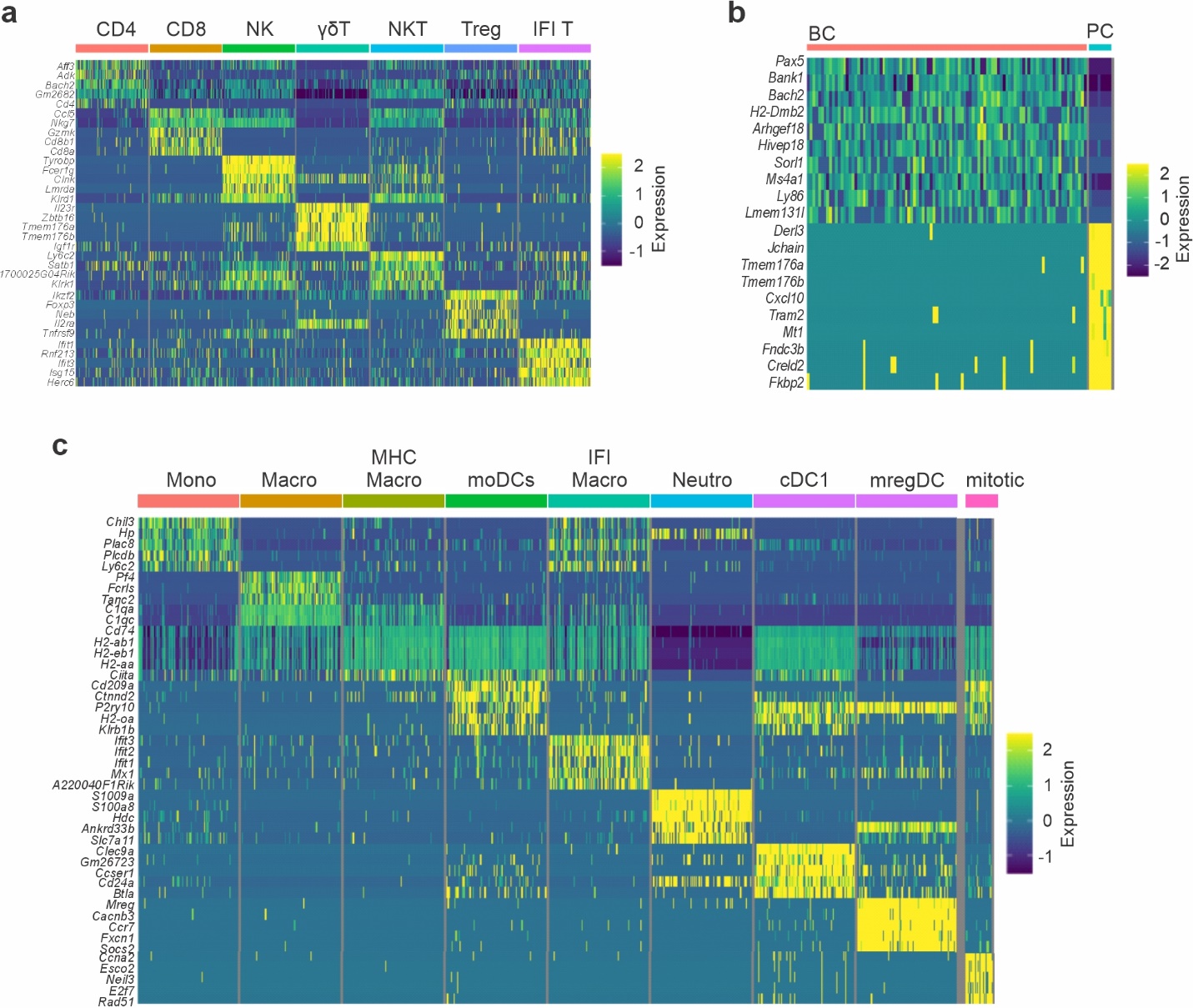

**Supplementary Figure 3.** **Delineation of immune cell subsets for T cell, B/Plasma cell, and myeloid clusters in CDK4/6-MEK inhibited *de novo* MPNSTs.** FindAllMarkers analysis of scRNA seq data was used to determine immune cell subsets within immune cell clusters in WT and PC-ko MPNSTs that had been treated for 4-6 days with CDK4/6-MEK inhibitors. Heatmaps of relative gene expression are listed for the: **a)** T cell cluster (Natural killer cells, NK; gamma-delta T cell, γδT; NKT cell, NKT; Regulatory T cell, Treg; and Interferon induced T cells, IFI T), **b)** B/Plasma cell cluster (B cell, BC; Plasma cell, PC), and **c)** the myeloid cluster (Monocyte, Mono; Macrophage, Macro; MHC-hi Macrophage, MHC Macro; monocytic dendritic cells, moDCs; Interferon induced macrophages, IFI Macro; Neutrophils, Neutro; classical DCs, cDC1; regulatory DCs, mregDC; proliferating myeloid cells, Mitotic).

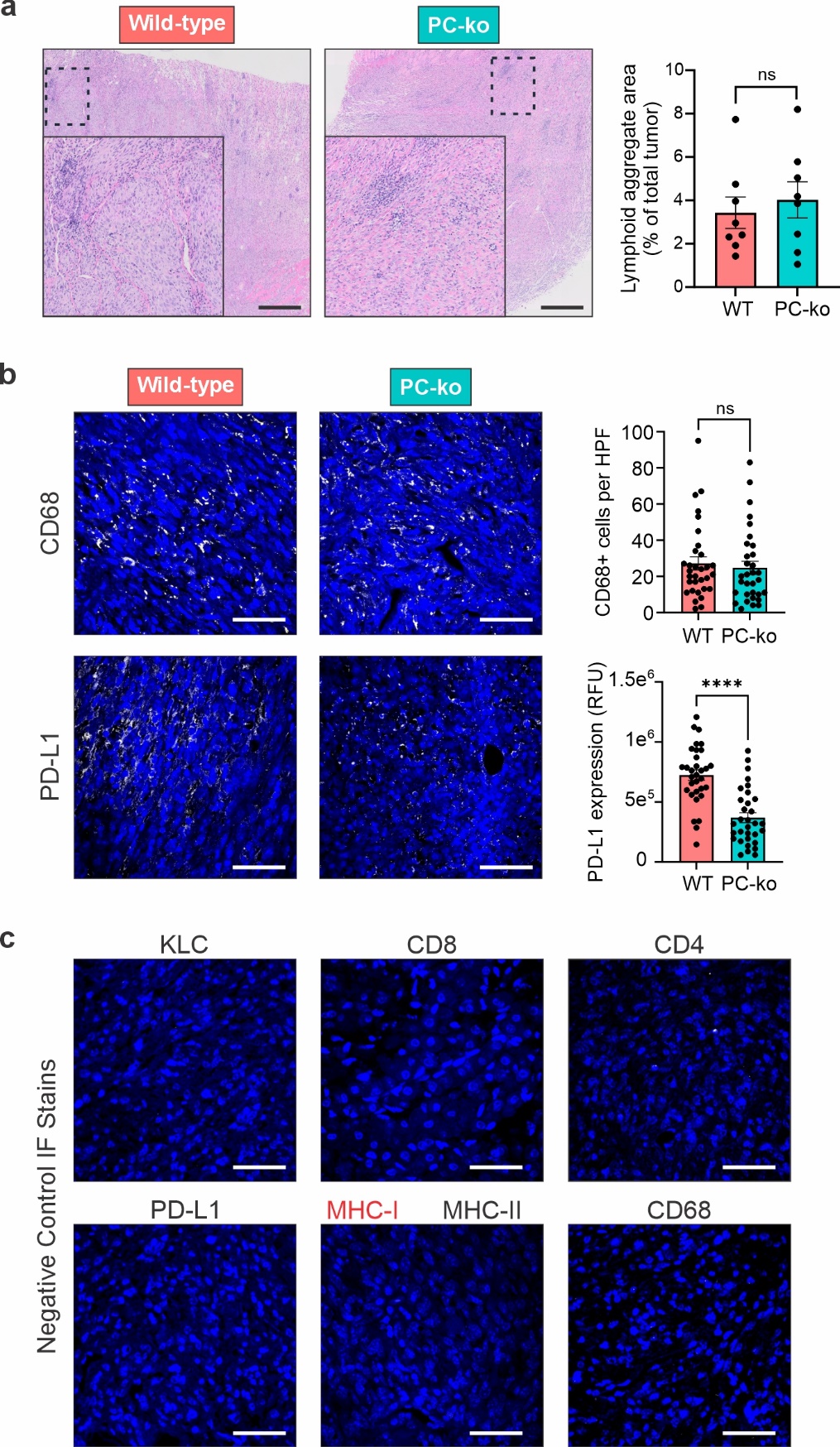
Supplementary Figure 4

**Supplementary Figure 4. Additional histopathological analyses of CDK4/6-MEK inhibited *de novo* MPNSTs. a)** Representative 10X images of H&E stained wild-type (WT) and PC-ko tumors with quantified area for lymphoid aggregates graphed to the right. Magnified region corresponds to the area in the black dashed box. Black scale bar represents 500 µM. Error bars are SEM. Statistical comparison done by Student’s t test; ns, not significant. **b)** Representative 40X images of IF staining for CD68 (top) and PD-L1 (bottom) in WT and PC-ko *de novo* MPNSTs with quantified data shown to the right**.** White scale bar represents 50 µM. Error bars are SEM. P value was obtained via Student’s t test (****, *P* < 0.0001); ns, not significant. **c)** Representative 40X images of IF staining results for negative control stains (no primary antibody) for each immunostaining protocol that employed distinct secondary antibodies. The target for each stain is indicated.

**Supplementary Table 1:** Gene-specific primer sequences used for quantitative RT-PCR

| **Gene** | **Forward Primer** | **Reverse Primer** |
| --- | --- | --- |
| *Cd4* | CTGATGTGGAAGGCAGAGAAG | GAGACCTGGGGTATCTTGAGG |
| *Foxp3* | CTCATGATAGTGCCTGTGTCCTCAA | AGGGCCAGCATAGGTGCAAG |
| *B2m* | AGACTGATACATACGCCTGCAG | CGAGGTTCAAATGAATCTTCAG |
| *H2-k1* | TCCATCCACTGTCTCCAACA | CTGGAGCCAGAGCATAGTCC |
| *H2-d1* | CATGGTGATCGTTGCTGTTC | CTGGAGCCAGAGCATAGTCC |
| *H2-Ab1* | ACCCAGCCAAGATCAAAGTGC | TGCTCCACGTGACAGGTGTAGA |
| *H2-Aa* | CTGACCACCATGCTCAGCTCT | TACTGGCCAATGTCTCCAGGAG |
| *Igkc* | GTGCCTCAGTCGTGTGCTTC | TGCTGCTCATGCTGTAGGTG |
| *Cxcl10* | GTGAGAATGAGGGCCATAGG | TTTTTGGCTAAACGCTTTCAT |
| *Cxcl9* | ATCTTCCTGGAGCAGTGTGG | AGTCCGGATCTAGGCAGGTT |
| *Gapdh* | GTTGTCTCCTGCGACTTCA | GGTGGTCCAGGGTTTCTTA |

**Supplementary Table 2:** Antibodies used for immunofluorescent detection of intratumoral immune cells and proteins.

| **Antibody** | **Vendor and Cat No.** | | **Dilution** | **Fluor** |
| --- | --- | --- | --- | --- |
| Kappa Light Chain | Abcam ab190484 | 1:50 | | 647 |
| CD8a | ThermoFisher MA1-10301 | 1:50 - amplified | | 647 |
| CD4-eFluor 660 | ThermoFisher 50-9766-82 | 1:10 | | none |
| MHC-I | ThermoFisher MA5011723 | 1:10 | | 568 |
| MHC-II | ThermoFisher 14-5321-82 | 1:100 | | 488 |
| CD68 | Abcam ab190484 | 1:100 | | 647 |
| PD-L1 | Proteintech 66248-1-Ig | 1:25 | | 568 |
